# Towards a holistic epidemiology of *Streptococcus agalactiae* using the BakRep repository

**DOI:** 10.64898/2026.03.02.709001

**Authors:** Linda Fenske, Oliver Schwengers, Alexander Goesmann

## Abstract

*Streptococcus agalactiae* is a versatile multi-host pathogen that can cause major neonatal disease in humans, as well as mastitis in dairy animals. Its ability to infect a wide range of hosts is largely driven by its high genomic plasticity and the acquisition of distinct accessory genes. The global population of *S. agalactiae* is characterized by multiple of capsular serotypes and clonal complexes that differ in their propensity to cause invasive disease, including hypervirulent CC17 (often serotype III) associated with neonatal meningitis, whereas CC1/CC19/CC23 are more often colonizing lineages. Although widely studied, most research is limited to particular regions or single outbreak events, offering only fragmented snapshots instead of a comprehensive global picture. To move beyond region- or outbreak-limited studies, this work has analyzed 37970 *S.agalactiae* genomes from BakRep, integrating serotypes, MLST, AMR genes, lineage-specific genes, and descriptive metadata to map current trends and identify potential gaps in public data. The dataset largely matched the known population structure with serotype III, Ia and V most common and stable serotype/clonal complex lineages (e.g. III-2/CC17, Ia/CC23, CC1/V), while also rising serotype diversity. Lineages differed in their accessory-gene profiles, with III-2/CC17 being enriched for virulence and adhesion genes, while other groups showed either greater genomic plasticity (mobile/phage genes) or niche specialization. AMR was widespread with very high tetracycline resistance (>80%), frequent MLSB resistance determinants, and emerging aminoglycoside resistance in some genomes. But overall it became evident that the associated metadata contained substantial gaps. Missing or incomplete information limits biological interpretation, underscoring that rigorously curated, structured metadata is essential for maximizing the value of ongoing sequencing efforts.

## Introduction

*Streptococcus agalactiae*, also known as Group B Streptococcus (GBS), is a Gram-positive bacterium, capable of causing infections in various host species. In human medicine, GBS is a common cause of neonatal infections, primarily transmitted from mother to newborn during childbirth (1). An estimated 20-30% of pregnant women are colonized with GBS, with around 50% of their newborns acquiring colonization and approximately 1% developing invasive disease (2,3). Asymptomatic GBS colonization is also commonly observed in both young adults (4) and elderly individuals (5). In the veterinary context, GBS is primarily known as a contagious udder pathogen linked to mastitis. While it is most commonly associated with mastitis in cattle, cases have also been reported in other dairy species, including goats (6,7), sheep (8,9), and camels (10). Although most published GBS research centers on human and bovine cases, GBS has also been detected in several other terrestrial (11,12) and aquatic species (13,14). The broad host range of GBS has been linked to its high genomic plasticity, enabling the integration of accessory genes into its chromosome, which allows strains to colonize diverse hosts and ecological niches, providing an evolutionary advantage (15).

The outer layer of GBS consists of a polysaccharide capsule, whose antigenic properties serve as the basis for further subclassification into currently ten major serotypes (Ia, Ib, II-IX) (16) and four subtypes of serotype III (III-1, III-2, III-3, III-4) (17). The prevalence and distribution of serotypes are known to vary depending on host species, levels of virulence or geographic region (18). Additionally, multilocus sequence typing (MLST) classifies GBS isolates into distinct sequence types (STs), which can be further grouped into clonal complexes (CCs). This can offer insights into the pathogenicity of isolates. For example, CC17, is known as a hyper-virulent cluster, increasingly associated with neonatal infections (19,20). Specifically the combination of serotype III and CC17 is considered to be responsible for the majority of neonatal meningitis cases (21) which is likely attributable to specific genes encoding secreted and surface proteins, which facilitate interactions between the bacterium and host cells (19). In contrast, CC1, CC19 and CC23 are common colonizers of pregnant women and appear well adapted to the vaginal mucosa, with relatively limited invasive potential in neonates (22,23).

As in general, antibiotic resistance (AMR) is also a growing concern in GBS strains. Intrapartum antibiotic prophylaxis (IAP) is administered to pregnant women based on risk factors or screening guidelines, making continuous monitoring of resistance rates essential (24). In mastitis cases, particularly in middle-income countries with higher GBS prevalence, widespread use of antimicrobials, including broad-spectrum and critically important antibiotics, contributes to antimicrobial resistance in multi-host pathogens, reducing treatment options for both animals and humans (25).

Numerous studies have analyzed GBS, though they are often restricted to specific geographic regions or isolated outbreak events. As a result, only limited subsets are analyzed rather than a more holistic overview of the entire population. In 2024, BakRep was released (26), featuring extensive genome characterizations based on 661405 uniformly processed assemblies of all short-read sequence data from the European Nucleotide Archive (ENA) up to November 2018 (27). Up to now, the collection of uniformly assembled data was expanded to 2440377 assemblies (28), which have since been characterized and made accessible through the BakRep web repository. With its vast collection of genomic information and associated metadata, this repository provides an ideal resource for obtaining a comprehensive overview of publicly available sequence data, enabling insights beyond individual studies.

In the present study, we conducted a comparative analysis of a total of 37970 GBS genomes contained in BakRep v2. We characterized all isolates by MLST, capsular serotypes, AMR genes, putative lineage-specific genes, and complemented this with metadata on geographic origin, host species, and disease patterns. This integrated analysis helps to evaluate current trends in the GBS study landscape, to highlight well-studied areas, and finally to identify understudied gaps with only scarce public data.

## Methods

### Export data from the BakRep repository

All genome data and associated metadata were retrieved from BakRep (https://bakrep.computational.bio/) (26). For this, the repository was searched for entries where the species exactly matches “Streptococcus agalactiae” (operator “eq”) and all search results were exported into a TSV file. Additional files provided by BakRep were downloaded via the BakRep CLI tool (https://github.com/ag-computational-bio/bakrep-cli) by providing the dataset ids of the identified GBS genomes.

### Data analysis

Capsular-serotypes were determined using KMA (v1.4.12) using the FASTA files as input (29). AMR genes were identified using AMRFinderPlus (v4.0.23) with default parameter settings (30). Bakta-generated GFF3 annotation files (31) were used as input for Panaroo (v1.5.0) (32). The resulting gene presence/absence matrix, combined with a binary trait file of selected phenotypes, was then used to run a pan-GWAS in scoary2 (v0.0.15) using default settings (33). The metadata retrieved from BakRep and all additional results have been analyzed using R (v4.4.2).

## Results

### Sequence statistics of GBS genomes

BakRep v2 contained 37970 genomes classified as *Streptococcus agalactiae*. The average estimated completeness of all GBS genomes was 99.94%, whereby only 98 (0.26%) genomes were below a completeness of 99%. The average estimated contamination was 0.49%, consisting of 139 (0.37%) genomes, which had a contamination lower than 50%, and a total of 529 (1.39%) genomes which had a contamination lower than 1% (Supplemental Figure 1).

The assembled contigs of all GBS genomes had total lengths ranging from 105259 to 12119634 base pairs (bp), with an average genome size of 2095432 bp. In the NCBI database, GBS genomes exceeding 2.6 Mbp are flagged as unusually large for this species. This size is exceeded by 206 genomes in BakRep. However, genomes exceeding this size threshold also exhibit a significantly higher rate of estimated contamination (data not shown). The GC content varied between 33.25% and 58.16%, averaging to 35.33%. The number of contigs per assembly ranged from 10 to 6016 with a mean of 51.59 (Figure 1).

**Figure 1:**
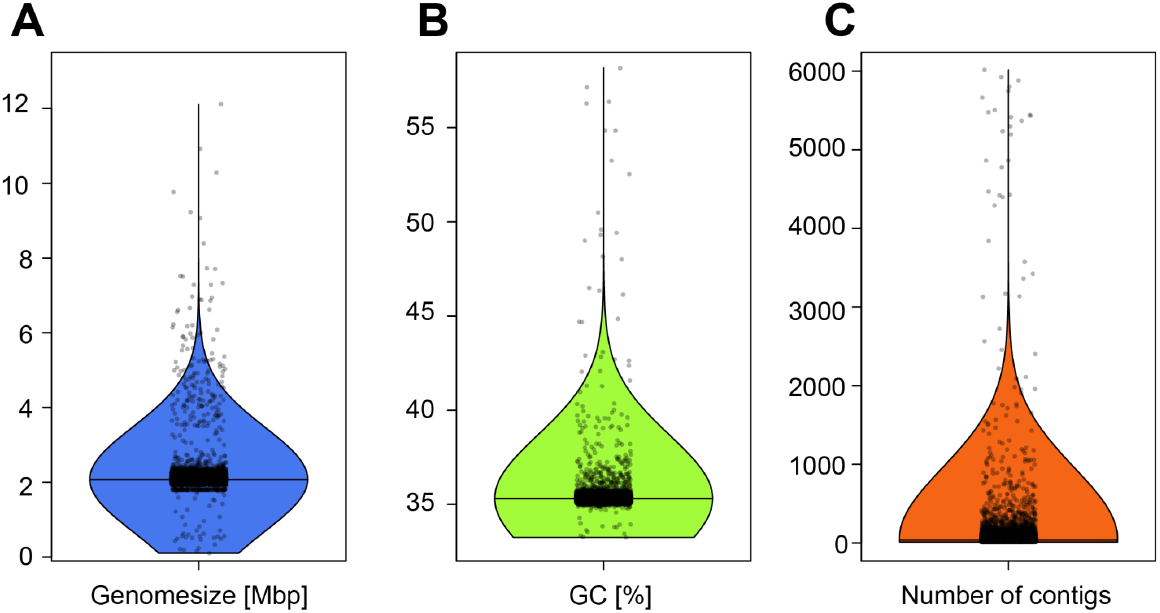
Comparison of genome size (**A**), GC content (**B**), and number of contigs (**C**) of all GBS genomes contained in BakRep v2 (n=37970). Individual dots represent single genomes, with horizontal jitter added to avoid overplotting. Horizontal black lines indicate medians; bold black bars represent interquartile ranges; vertical black lines represent outliers.

### Metadata of GBS genomes

The metadata associated with the GBS genomes in BakRep v2 indicated that the genomes originate from 51 countries across six continents. The majority of genomes derive from North America (49.73%; n=18884), with the United States alone accounting for 48.59% (n=18451) of all genomes. No isolation region was provided for 14747 (38.84%) genomes (Figure 2).

**Figure 2:**
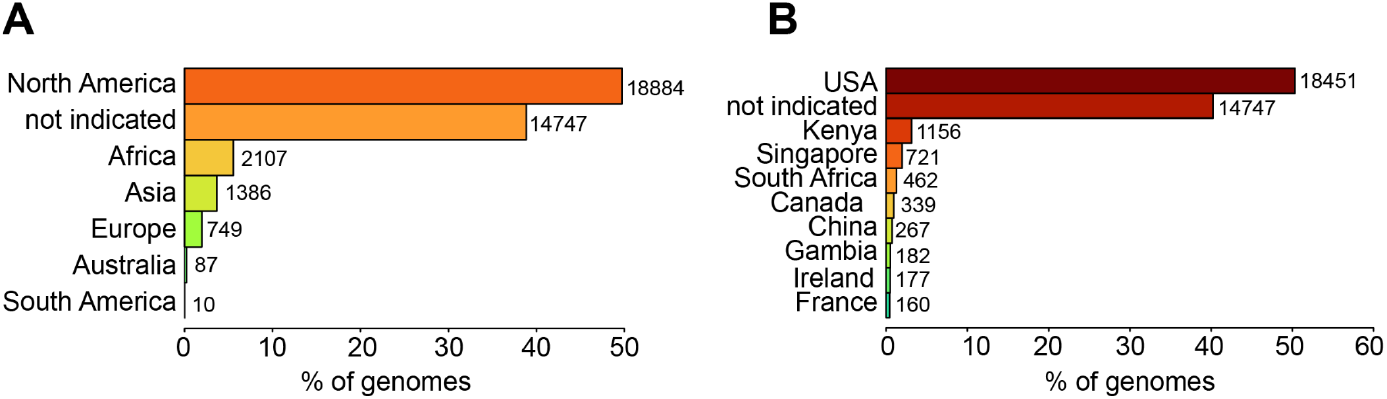
Distribution of GBS genomes contained in BakRep v2 across different continents (**A**) and the ten most common countries (**B**). The total number of genomes for each country and continent, respectively, is displayed right to the bars.

The majority of genomes were declared as human isolates (58.41%; n=22179), followed by 61 bovine-derived genomes (0.16%) and 48 genomes obtained from different fish species (0.12%). Three (0.01%) genomes were listed as isolates from dogs, while individual (0.004%, each) genomes were obtained from a cat, a llama, and a crocodile, respectively. A host species was not provided for 41.28% (n=15675) of genomes. For 98.35% (n=37342) of the genomes, no information on disease status was provided; 0.88% (n=334) were reported as diseased, 0.60% (n=226) as healthy, and 0.16% (n=60) as carriage isolates. The collection date spans from 1972 to 2024. Regarding the yearly distribution of isolates, a gradual increase is observed over time. Between 1972 and 2005, genome counts remained very low, but with a noticeable increase from 2006 onwards. The majority of genomes were collected between 2015 and 2022, with a slight decline observed after 2020 and very few genomes available for 2023 and 2024 (Supplemental Figure 2). For 39.05% (n=14829) of genomes no information for the isolation year was provided.

### Serotype, Sequence Type and Clonal complex distribution

Some genomes had capsular serotype information provided in the associated metadata. However, since the methodology used for serotype determination was unclear and 98.02% (n=37629) genomes lacked serotyping, capsular serotypes were *in silico* reanalyzed for all genomes. The analysis of the serotype distribution revealed that most GBS genomes belonged to one of the nine predominant GBS serotypes plus all four subtypes of serotype III. Only 97 (0.26%) genomes were non-typable. Serotype III was the most common, accounting for 24.41% (n=9270) of all genomes, followed by serotype Ia at 21.07% (n=8000), and serotype V at 16.25% (n=6172) (Figure 3A).

**Figure 3:**
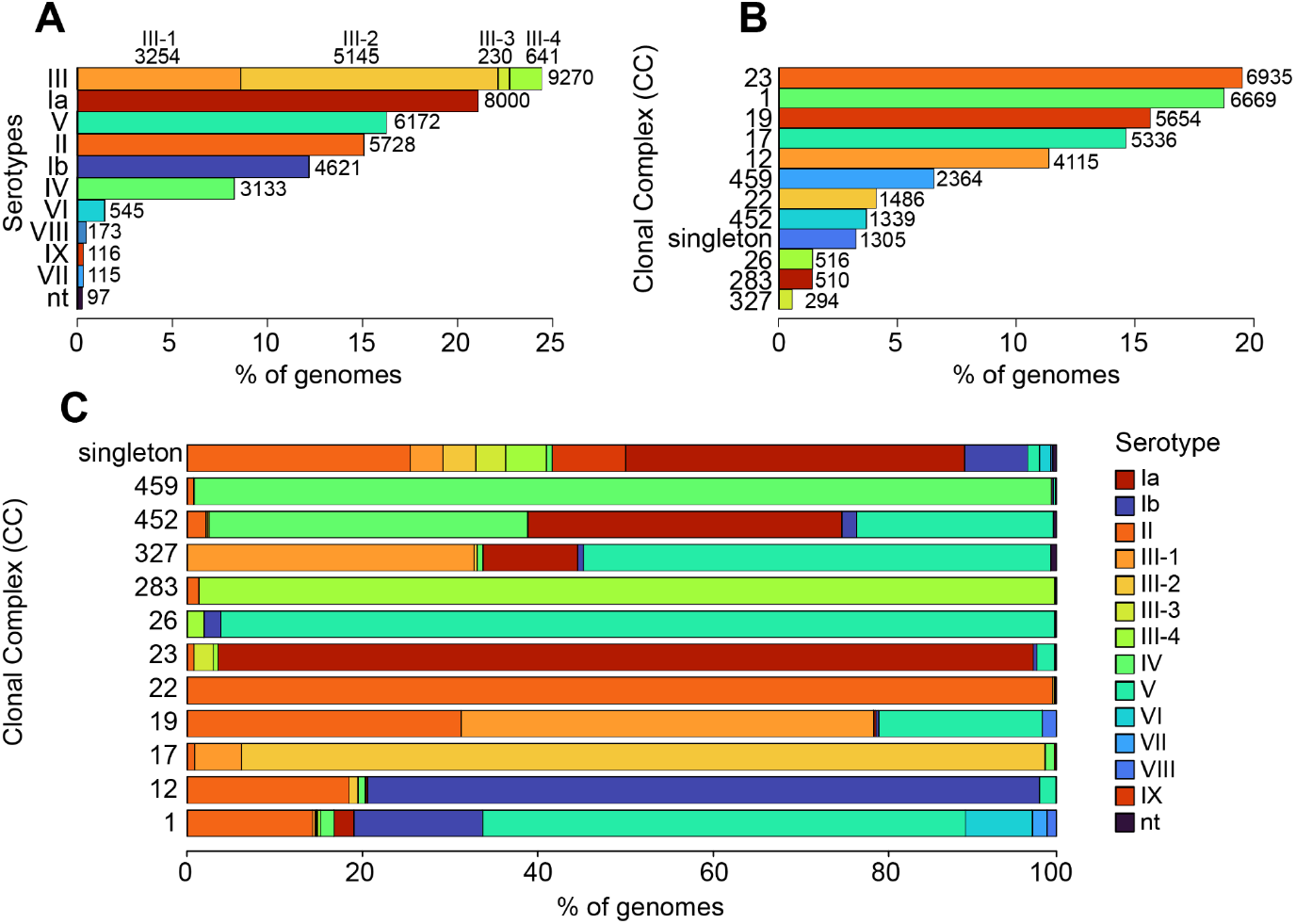
Serotype **(A)** and CC **(B)** distribution, and serotype allocation **(C)** among the CCs of all GBS genomes contained in BakRep v2. The total number of genomes for each serotype and CC, respectively, is displayed right to the bars.

MLST analysis identified 1015 unique STs, which covers 40.39% of the STs currently listed in PubMLST. ST23 was most prevalent at 14.56% (n=5530) of all genomes, followed closely by ST1 at 14.31% (n=5434) and ST17 at 11.62% (n=4414). The STs could be grouped into eleven CCs. The main CCs among all GBS genomes were CC23 (18.99%; n=6935), CC1 (18.25%; n=6669), and CC19 (15.48%; n=5654) (Figure 3B).

The analysis of the associations between CCs and serotypes revealed several connections. CC23 was strongly associated with serotype Ia, which accounted for 93.73% (n=6500) of those genomes. CC17 was mainly associated with serotype III-2, which accounted for 92.38% (n=4929) of those genomes. CC459 was nearly exclusively linked to serotype IV (98.60%; 2331), while CC22 was similarly linked to serotype II (99.53%; n=1479). CC1 showed a predominance of serotype V (55.56%; n=3705), followed by substantial proportions of serotypes Ib (14.78%; n=986), II (14.42%; n= 962) and VI (7.67%; n=512), indicating a more moderate serotype diversity within this lineage. CC19 presented an even more heterogeneous serotype distribution, with serotype III-1 (47.40%; n=2680), II (31.52%; n=1782) and V (18.84%; n= 1065) co-occurring (Figure 3C). Although most serotypes were associated with multiple STs, serotype IX was found predominantly in ST130 (93.10%; n=108). Exceptions included a single genome each in ST1208 and ST1216, both single-locus variants (SLVs) of ST130, as well as one occurrence in ST24, which belongs to CC452. Similarly, serotype VI was largely restricted to CC1, except for two occurrences in CC12 and 16 occurrences in ST889. Serotype VII was likewise primarily associated with CC1, except for one occurrence in ST103 and one occurrence in ST1438. All detailed numbers are listed in Supplementary Table S1.

### Continental and temporal variation in serotypes and clonal complexes

Serotype and CC distributions showed geographic variation. In North America, serotypes Ia (20.74%; n=3917), III (18.43%; n=3480), distributed across all III subtypes, and II (17.90%; n=3380) were most frequently observed (Figure 4A). In contrast, serotype III (42.76%; n=901) predominated in Africa, particularly subtypes III-1 and III-2 (Figure 4B). Asia showed a distinct pattern, with serotype III-4 (28.14%; n=390) being most prevalent, a subtype that was rare in other regions (Figure 4C). In Europe, serotypes III-2 (24.03%; n=180), Ia (20.83%; n=156), and II (18.02%; n=135) were among the most common, whereas in Australia, serotype Ib (29.89%; n=26) was the most frequently identified (Figure 4D+E). Regarding clonal complexes, CC1 (20.68%; n=3763) and CC23 (19.78%; n=3599) were dominant in North America. In Africa and Europe, CC17 (30.75%; n=643 and 32.86%; n=230) was the most prevalent, while Asia was characterized by a high proportion of CC283 (28.11%; n=377) and CC1 (27.89%; n=374). Notably, both serotype III-4 and CC283 were found almost exclusively in Asia.

**Figure 4:**
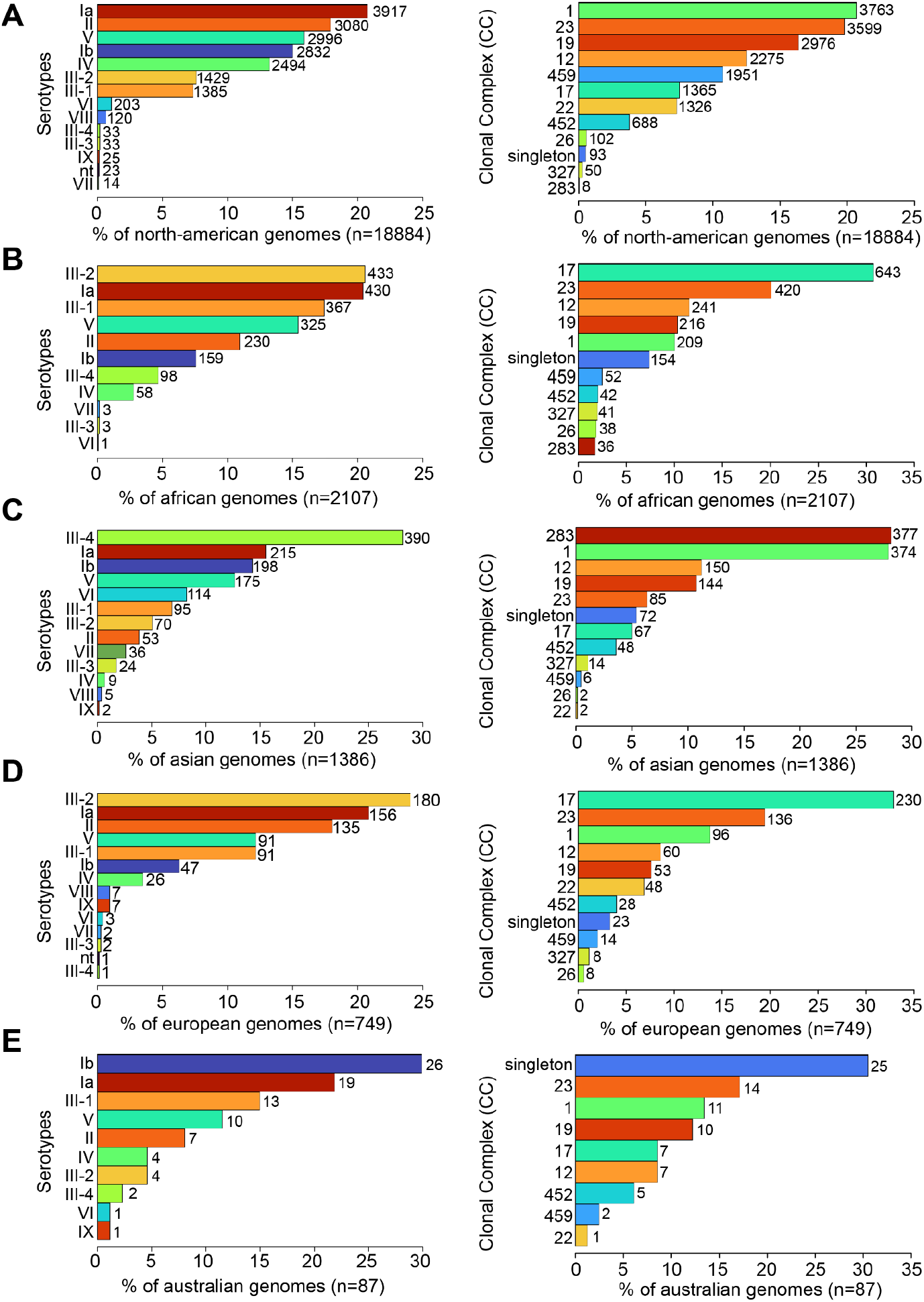
Serotype (left) and CC (right) distribution by continent. (**A**) North-American genomes, (**B**) African genomes, (**C**) Asian genomes, (**D**) European genomes and (**E**) Australian genomes. The total number of genomes is displayed right to the bars.

Examining the temporal distribution of serotypes revealed that serotype Ia has remained consistently prevalent over the years, comparable to serotype V, which, however, shows greater fluctuations. Serotype Ib and II showed a gradual increase, whereas III-1 appeared to have slightly declined. In contrast, serotypes III-3, VIII and IX have consistently remained at very low frequencies (Figure 5A). The temporal distribution of CCs showed an increase in diversity and abundance over time. Prior to the early 1990s, CCs were largely restricted to CC1. From the mid-1990s onwards, additional CCs such as CC17, CC19, CC23 and CC283 emerged, although at relatively low frequencies. From 2004 onwards there was a concurrent presence and rapid increase of multiple dominant lineages, including CC1, CC12, CC17, CC19 and CC23 (Figure 5B). Detailed numbers for each year are listed in Supplementary Table S2.

**Figure 5:**
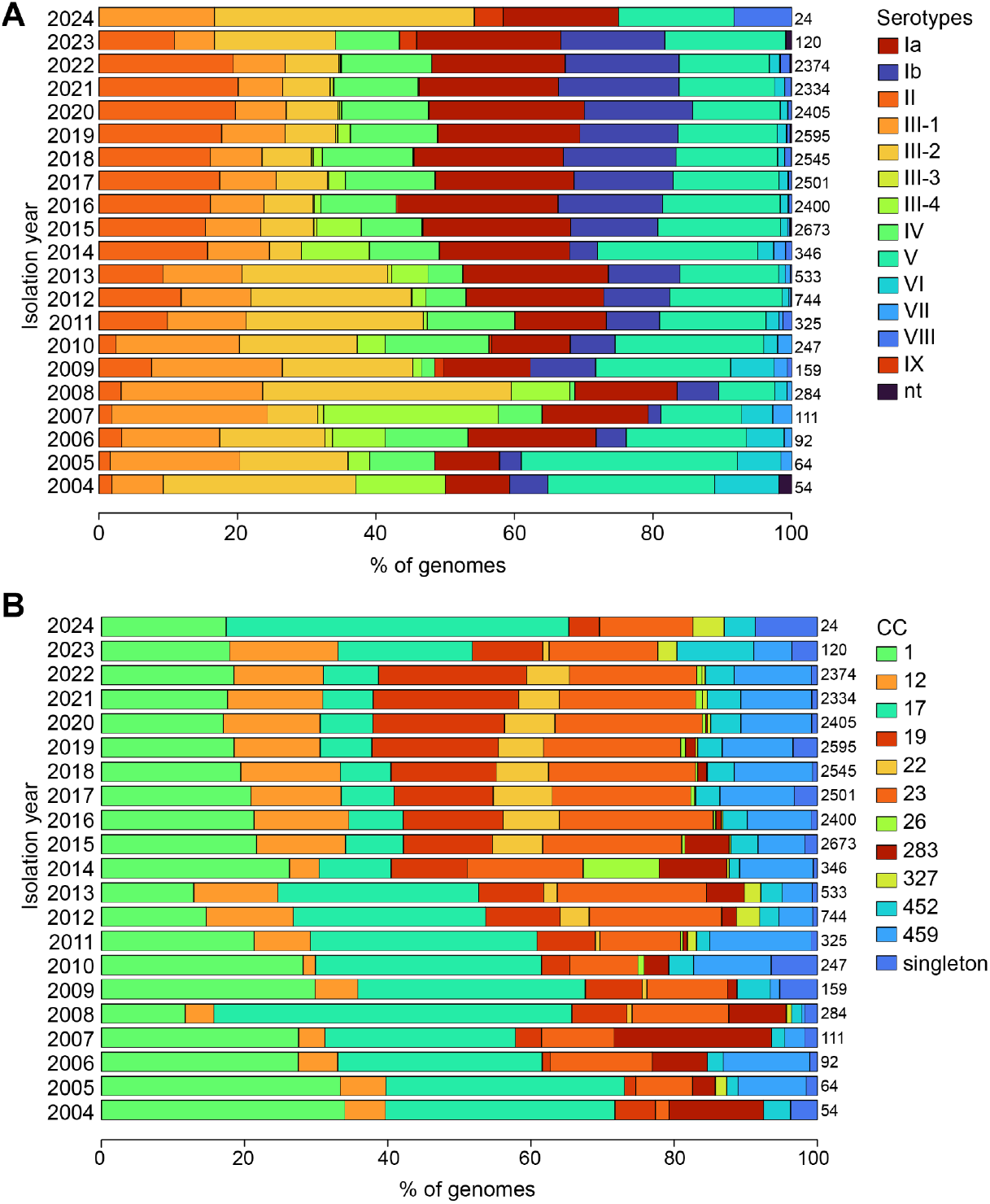
Serotype **(A)** and CC **(B)** distribution per year. Displayed are the years 2004-2024. The total number of genomes for each isolation year is displayed right to the bars.

### Resistome of GBS genomes

A total of 88.10% (n=33452) GBS genomes harbored at least one AMR gene. In total 132 known AMR gene determinants were identified conferring to resistance against 24 drug classes. Tetracycline resistance genes (a total of 16 different) were the most prevalent AMR determinants, observed in 84.24% (n=31985) of all genomes. Of these, the most common genes were *tet(M)* identified in 76.99% (n=29232) of genomes and *tet(O)* identified in 8.42% (n=3196) of genomes. The second most common were genes conferring for macrolide-lincosamide-streptogramin B resistance (MLSB) present in 17.66% (n=6707) of all GBS genomes. Of that, most common were *erm(B)* observed in 16.38% (n=6220) of all genomes, *erm(A)* observed in 13.11% (n=4976) of genomes and *erm(T)* observed in 1.3% (n=492) of genomes. 4.15% (n=1574) of all genomes carried aminoglycoside resistance genes. Only 0.14% (n=55) of all genomes carried beta-lactam resistance genes.

More than one AMR gene hit was found for 41.34% (n=15695) of the genomes, including 447 genomes with more than five hits and 11 genomes with more than ten hits. In total, 40.57% (n=15403) of genomes carried resistance genes for multiple drug classes, and 2.89% (n=1098) harbored genes for three or more classes, classifying them as multidrug-resistant. Notably, in one genome, resistance genes against as many as seven different drug classes were detected (ID: SAMEA112769107).

Since erythromycin and tetracycline resistance genes are often located on the same transposon, we examined their co-occurrence to determine in how many genomes both genes were present simultaneously and how much only contained one gene variant individually. The combination of a erythromycin and tetracycline resistance gene was present in 27.62% (n=10485) of genomes, while a tetracycline resistance gene alone was detected in 56.62% (n=21500) and an erythromycin resistance gene alone in 2.92% (n=1109). The combination of *erm(B)* and *tet(M)* was the most common combination at 1.1% (n=4180) of all genomes followed by the combination of *erm(A)* and *tet(M)* at 1.04% (n=3963), and the combination of *erm(B)* and *tet(O)* at 0.58% (n=2209). The combination of *tet(M)* and *mef(A*) was detected in 7.86% (n=2979) of all genomes and the combination of *tet(O*) and *mef(A)* was present in 0.48% (n=183) of all genomes.

Furthermore, macrolide resistance in streptococci is mediated by an efflux system, with *mef(A)* encoding the transmembrane channel and *msr(D)* the ATP-binding domains (34). We therefore analyzed the presence of *mef(A)* and *msr(D)* across all genomes, identifying both genes in 8.52% (n=3236) of genomes, while *mef(A)* alone was found in 0.04% (n=16) and *msr(D)* alone in 0.01% (n=4). Detailed numbers for all AMR genes and classes are listed in Supplemental Table 3.

### Serotype and CC associated resistome patterns

Among the eleven CCs, CC1 harbored the highest number of distinct AMR genes (n=72). *Tet(M)* was the most prevalent gene across all CCs, with the exception of CC459. CC459 stood out with *erm(A)* as its most common gene, showing a markedly higher frequency compared to other CCs (Figure 6F). Aminoglycoside resistance genes, such as *ant(6)-Ia* and *aph(3’)-IIIa*, were detected in all CCs, with the highest abundance observed in CC17 (Figure 6D). CC17 also harbored the highest number of hits for the trimethoprim resistance gene *dfrF*. Several Vancomycin resistance genes were exclusively found in CC1 (*vanS-G, vanY-G1, vanR-G, vanU-G*). The Macrolide and Strepogramin resistance gene *msr(D)* was by far the most common found in CC23 (Figure 6A). Serotype II showed the highest AMR gene diversity, encompassing 72 unique resistance determinants, with *tet(M)* being universally present. Notably, in serotype VIII, *tet(O)* was more prevalent than *tet(M)*. Similar to CC459, serotype IV was characterized by a high prevalence of *erm(A)*. In contrast, serotype Ia exhibited fewer *erm(A)* and *erm(B)* hits but a higher occurrence of *mef(A)* and *msr(D)* (Figure 7). The highest proportion of genomes without any AMR hits was observed in CC283 (58.82%), followed by CC452 (31.76%). Among serotypes, serotype IX showed the largest fraction without detected AMR genes (95.69%) followed by serotypes VIII (78.03%), VI (61.16%), and III-4 (51.34%).

**Figure 6:**
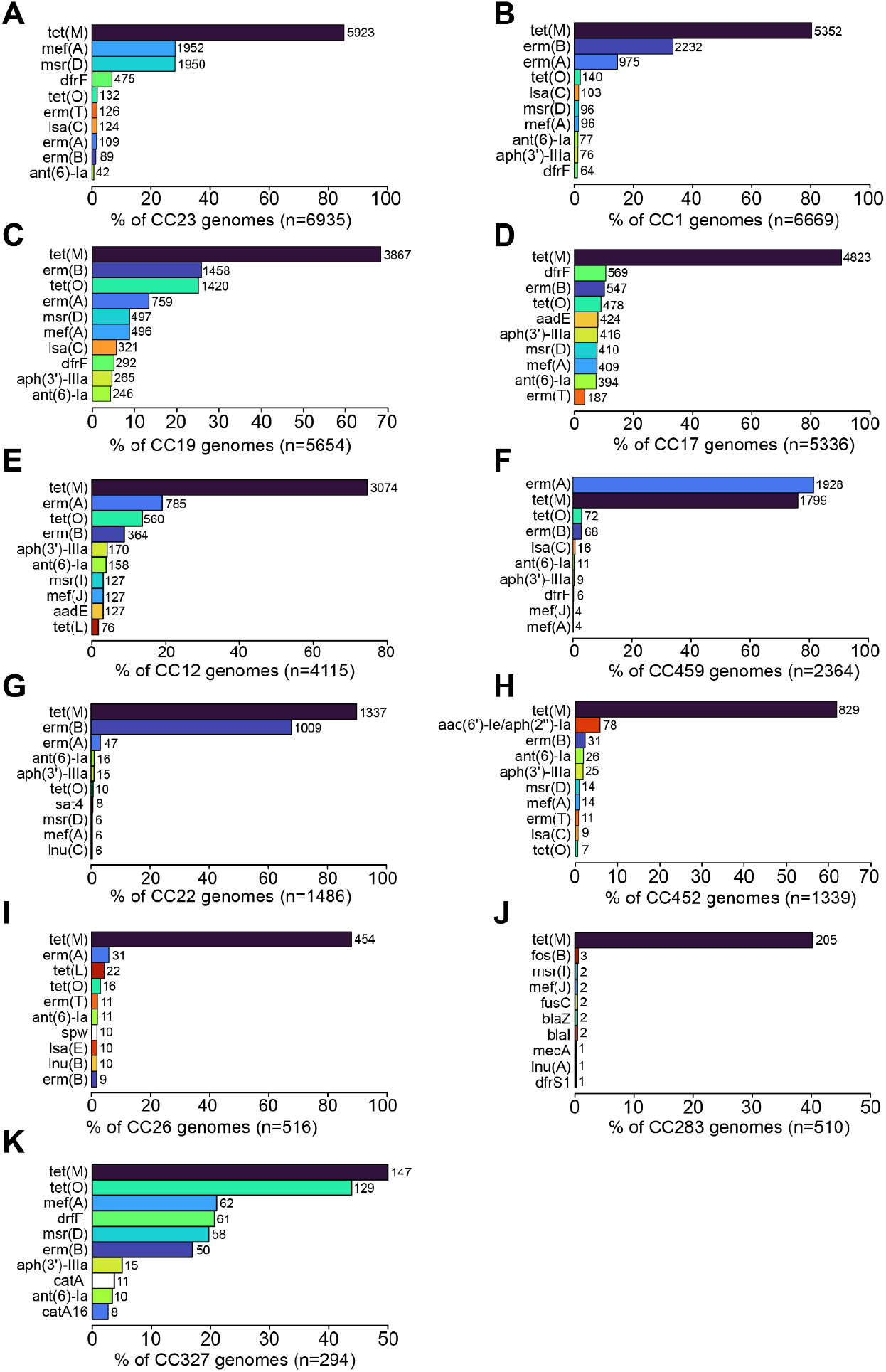
Distribution of the ten most prevalent AMR genes per CC. (**A**) CC23, (**B**) CC1, (**C**) CC19, (**D**) CC17, (**E**) CC12, (**F**) CC459, (**G**) CC22, (**H**) CC452, (**I**) CC26, (**J**) CC283, (**K**) CC327. The total number of genomes for each AMR gene is displayed right to the bars.

**Figure 7:**
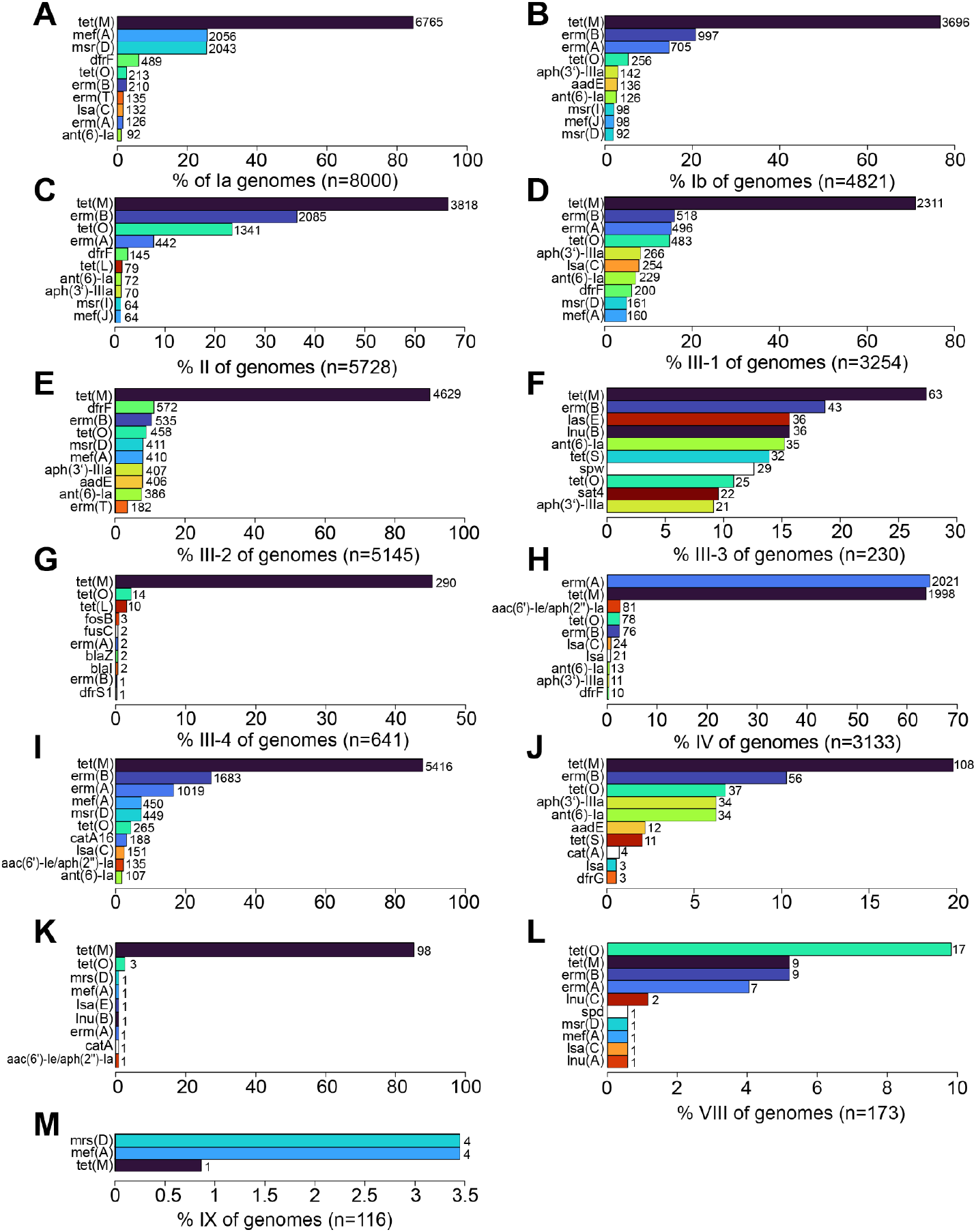
Distribution of the ten most prevalent AMR genes per serotype. (**A**) Ia, (**B**) Ib, (**C**) III-1, (**D**) III-2, (**E**) III-3, (**F**) III-3, (**G**) III-4, (**H**) IV, (**I**) V, (**J**) VI, (**K**) VII, (**L**) VIII, (**M**) IX. The total number of genomes for each amr gene is displayed right to the bars.

### Regional resistome patterns

Additionally, the geographical distribution of AMR genes was analyzed. *Tet(M)* emerged as the most widespread resistance gene, detected across all continents. North America exhibited the highest diversity and prevalence of AMR genes (n=59), especially those conferring resistance to macrolides and tetracyclines (Figure 8A). In Africa, several less common resistance genes were identified, including *dfrF, aac(6’)-Ie/aph(2’’)-Ia, msr(I), mef(J)*, and *catA16* (Figure 8B). The resistance gene profiles in Asia and Europe were largely similar, albeit with slightly lower overall prevalence (Figure 8C+D).

**Figure 8:**
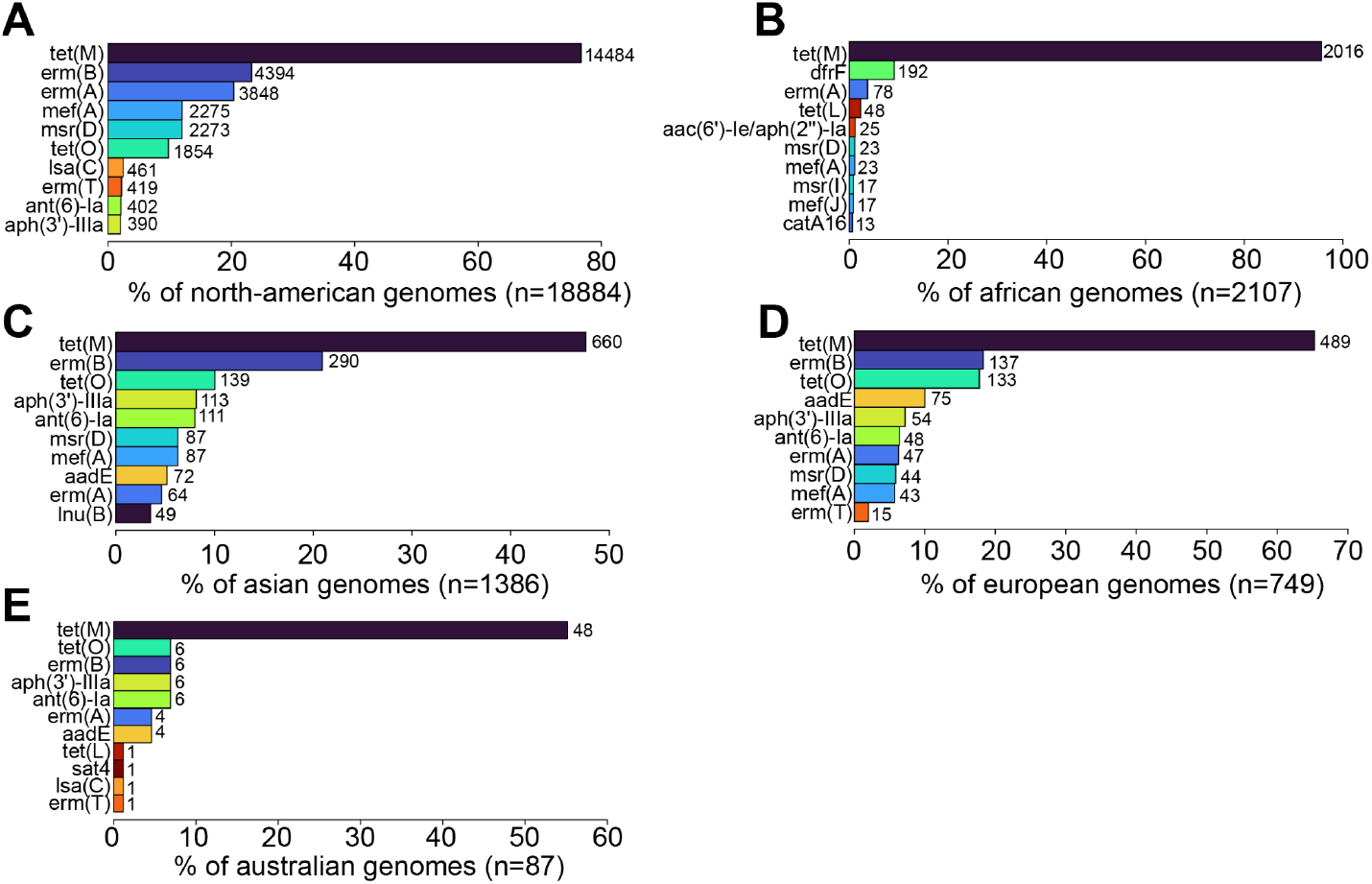
Distribution of the ten most prevalent AMR genes per continent. (**A**) North-America, (**B**) Africa, (**C**) Asia, (**D**) Europe, (**E**) Australia. The total number of genomes for each amr gene is displayed right to the bars.

### Temporal resistome variations

AMR genes were largely absent in isolates collected prior to the 1990s, with only sporadic occurrences of certain tetracycline and macrolide resistance genes appearing in the late 1980s. *Tet(M)* remained the most prevalent gene throughout the past two decades but since 2008, the prevalence of *tet(O)* has also risen markedly. MLSB genes like *erm(A)* and *erm(B)* became more frequent from 2000 onward, showing a steady increase. For aminoglycoside resistance genes, there was no consistent upward trend over time, instead only sporadic peaks are observed. Over all, AMR genes have shown a clear upward trend, with a gradual increase in diversity and frequency over the last 20 years (Figure 9). Detailed numbers for each year are listed in Supplementary Table S2.

**Figure 9:**
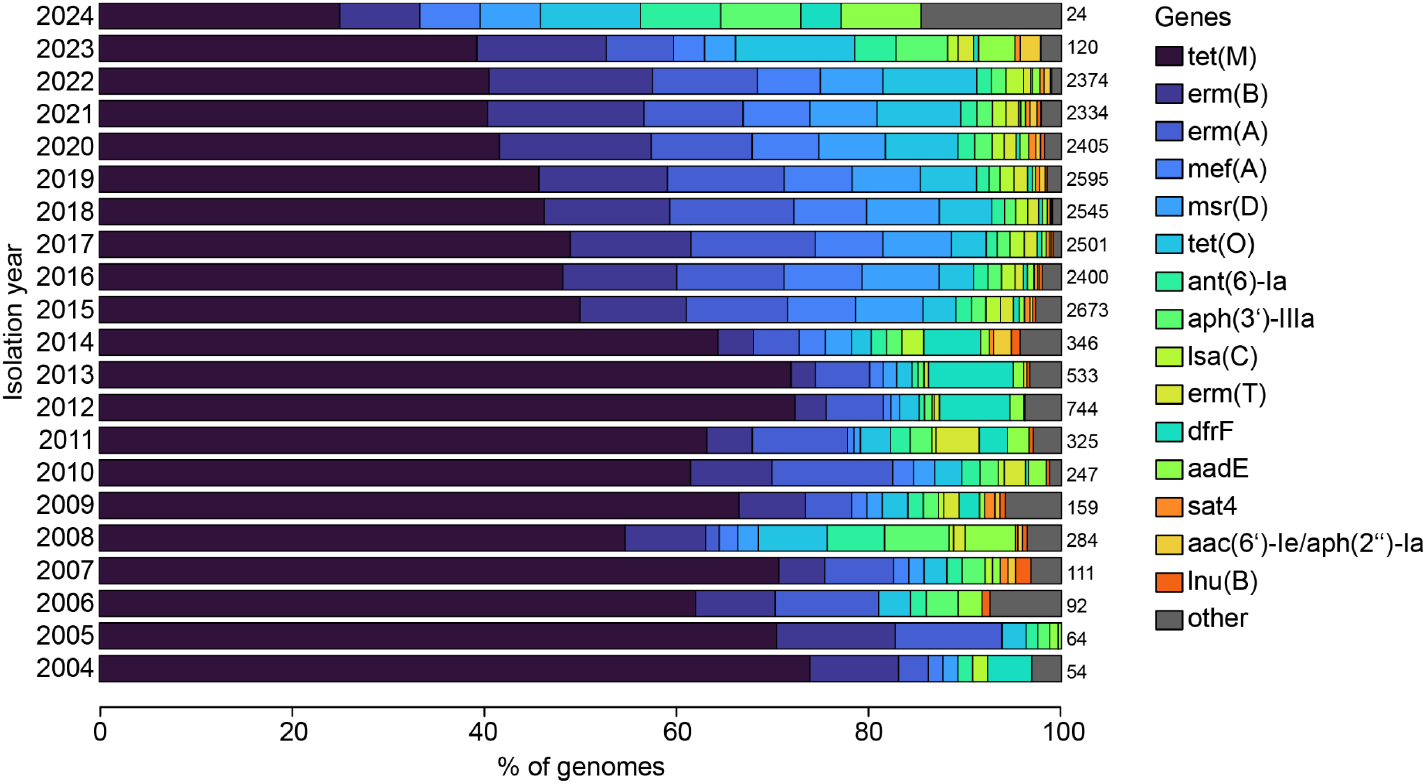
AMR gene distribution per year. Displayed are the years 2004-2024. The total number of genomes for each isolation year is displayed right to the bars.

### Pan genome analysis and genome-wide association investigation

Gene-content analysis of the 37970 GBS genomes identified a pan genome of 274610 genes and a core genome of 1398 genes. The distribution of genes displayed that most genes occurred in only a small fraction of genomes, with more than 99% of genes present in fewer than 15% of isolates (cloud genes). Accordingly, we filtered the gene presence-absence matrix out of Panaroo to retain genes present in more than 0.25% and less than 95% of genomes, resulting in 4101 remaining genes (Figure 10). The phylogenetic analysis done by scoary2 showed, although the genomes cluster partly according to serotype and CC, there are also areas with very diverse distribution (Figure 11).

**Figure 10:**
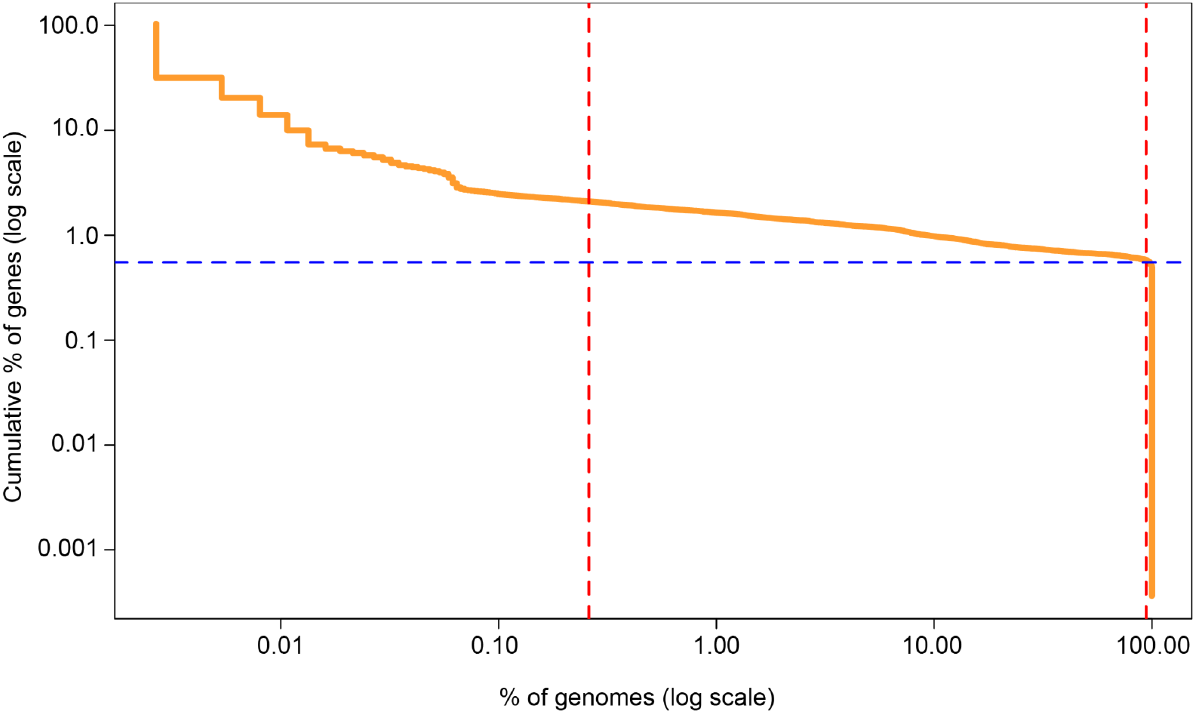
Pan-genome gene frequency distribution. The x-axis represents the percentage of genomes containing a given gene, and the y-axis represents the cumulative percentage of genes, both on logarithmic scales. Vertical red dashed lines indicate the lower and upper frequency cutoffs (0.25% and 95% of genomes), while the blue dashed horizontal line marks the core genome.

**Figure 11:**
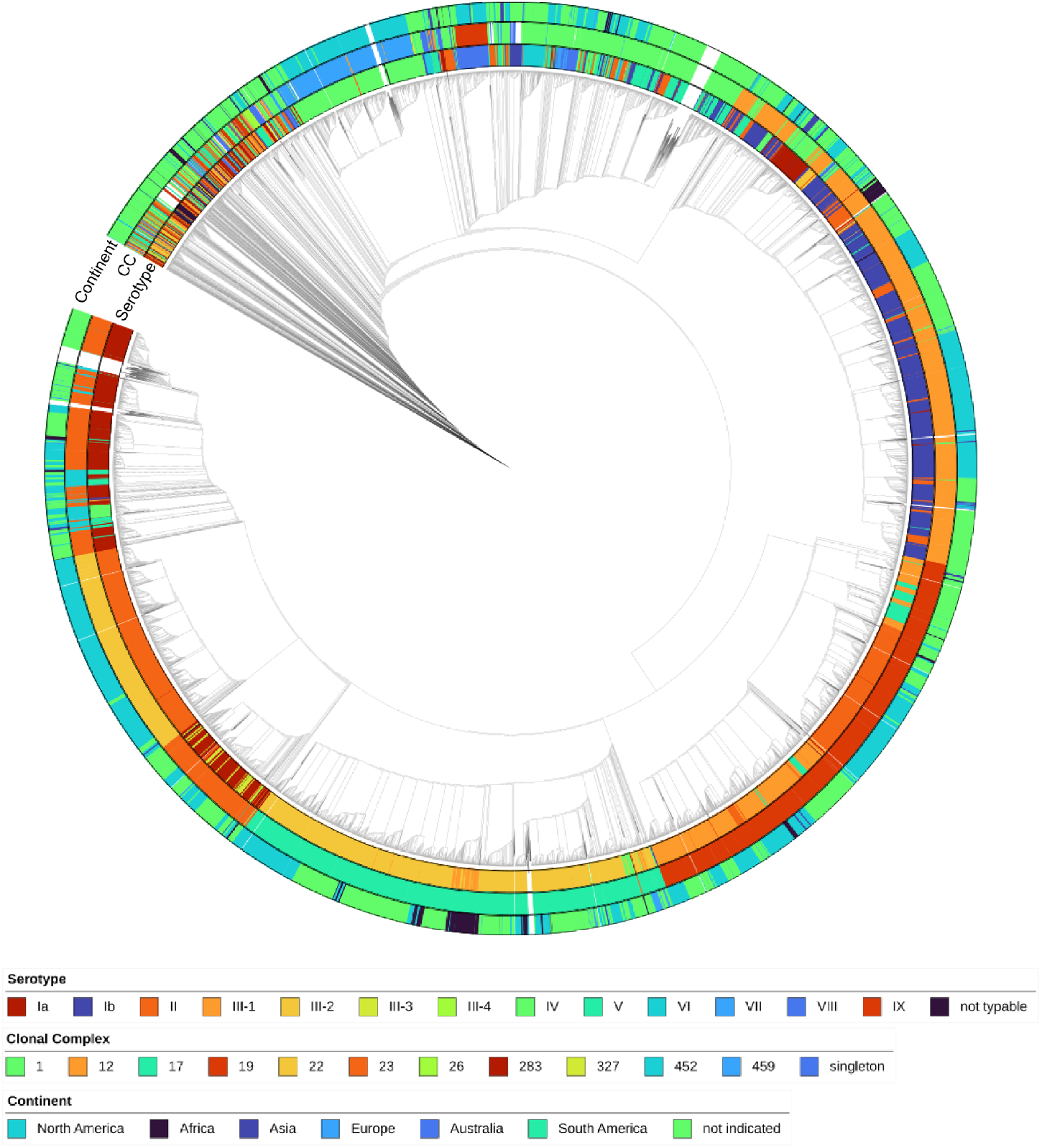
Phylogenetic tree of 37970 GBS genomes. The tree was constructed with scoary2. Colored blocks show serotype, clonal complex and continent of isolation. Annotation of the tree was done using iTol.

Scoary2 analyses were performed on the five most common serotype/clonal-complex combinations (Ia/23, II/22, III-2/17, IV/459, V/1) to identify genes predominantly associated with each combination. Multiple genes showed positive and negative associations with each group. Specifically, 60, 54, 41, 86, and 71 genes were positively associated with Ia/23, II/22, III-2/17, IV/459, and V/1, respectively, while 31, 12, 17, and 1 genes were negatively associated, with more than 70% specificity and sensitivity. Notably, group III-2/17 contained the highest number of positively associated genes with a specificity and sensitivity greater than 90% (n=30) followed by group II/22 (n=17). Group Ia/23 harbored several mobile genetic elements, numerous phage-associated genes and loci involved in metal homeostasis and heavy-metal resistance. In contrast, groups II/22 and III-2/17 encoded more functionally specialized genes. II/22 was enriched for genes related to metal uptake, including siderophore biosynthesis, whereas III-2/17 showed a marked accumulation of virulence and adhesion determinants, particularly components of the accessory secretion system (SecA2/SecY2 and Asp proteins). Groups IV/459 and V/1 were dominated by phage-associated genes. Table 1 lists the top 3 positive and negative hits per group. All other genes are listed in Supplementary Table S4.

**Table 1:** Scoary2 results are reported for the most significant positively (+) and negatively (-) associated genes of the five groups ((Ia/23, II/22, III-2/17, IV/459, V/1). The top 3 hits positive and negative are displayed. **SE**: Sensitivity of using the presence/absence of this gene as a diagnostic test to determine trait-positivity. **SP**: Specificity of using the absence/precence of this gene as a diagnostic test to determine trait-negativity. **P-value**: The *p*-value of the post-hoc permutation for the best gene.

| Phenotype | pos(+)<br>/neg (-) | Gene | Annotation | SE | SP | *p*-value |
| --- | --- | --- | --- | --- | --- | --- |
| **Ia/23**<br>**(n=6497)** | + | group_40999 | Phosphoribosylaminoimidazole<br>carboxylase ATPase subunit | 100.0 | 91.71 | 0.0020 |
|  | + | group_51961 | DUF6287 | 100.0 | 91.69 | 0.0020 |
|  | + | *purN* | Phosphoribosylglycinamide<br>formyltransferase | 99.98 | 88.82 | 0.0020 |
|  | - | group_9603 | GNAT family<br>N-acetyltransferase | 99.45 | 94.18 | 0.0020 |
|  | - | *aprE* | S8 family serine peptidase | 99.46 | 85.12 | 0.0020 |
|  | - | group_11313 | hypothetical protein | 99.86 | 73.96 | 0.0020 |
| **II/22** | + | group_41231 | DUF6261 | 100.0 | 98.61 | 0.0020 |
|  | + | group_52008 | Membrane protein | 100.0 | 98.60 | 0.0020 |
|  | + | group_51962 | Phage protein | 100.0 | 98.60 | 0.0020 |
|  | - | group_6272 | Cingulin | 100.0 | 81.76 | 0.0020 |
|  | - | group_23506 | Tandem five-TM protein | 100.0 | 73.78 | 0.0020 |
|  | - | group_19754 | KTSC domain-containing<br>protein | 100.0 | 73.67 | 0.0020 |
| **III-2/17** | + | group_6225 | Cell wall surface anchor protein | 99.98 | 90.46 | 0.0020 |
|  | + | *spaA*~~~inIB | Sortase | 99.96 | 90.31 | 0.0020 |
|  | + | group_10458 | S8 family serine peptidase | 99.94 | 96.21 | 0.0020 |
|  | - | group_41244 | HIT family protein | 99.96 | 74.90 | 0.0020 |
|  | - | group_9833 | Lipoprotein | 99.88 | 87.97 | 0.0020 |
|  | - | group_9933 | TIGR04197 family type VII secretion effector | 99.88 | 71.98 | 0.0020 |
| **IV/459** | + | group_9154 | Integrase | 99.96 | 98.95 | 0.0020 |
|  | + | *padR* | Transcriptional regulator | 99.96 | 98.95 | 0.0020 |
|  | + | group_41269 | N-acetyltransferase | 99.96 | 98.95 | 0.0020 |
|  | - | group_11745 | Relaxase | 99.96 | 77.27 | 0.0020 |
| **V/1** | + | neuA~~~wca<br>A~~~cpsJ | Glycosyltransferase (Capsular polysaccharide biosynthesis protein) | 98.49 | 91.80 | 0.0020 |
|  | + | group_25544 | Phage protein | 98.22 | 76.39 | 0.0020 |
|  | + | group_24863 | Phage protein | 98.22 | 76.20 | 0.0020 |

Other works reported the *scpB-lmb* transposon mainly associated with the human host (35,36). Because most genomes were decelerated as human isolates, we examined how frequently these genes occurred in the dataset. The *scpB* gene was detected in 99.91% (n=37934) of genomes and *lmb* in 94.85% (n=36013). However, 55 distinct *scpB* hits were found by Panaroo, whereas *lmb* was represented by a single variant.

## Discussion

*Streptococcus agalactiae* is a known multi-host pathogen, primarily linked to infections in humans and cattle. Although often studied, most investigations focus on specific regions or outbreaks, leaving a lack of global perspective on GBS diversity. BakRep, with its data size and integration of genomic data and metadata, offers an opportunity to examine study trends on a broader scale and supports more holistic analyses of the GBS population as a whole. Based on this genome repository, in total 37970 GBS genomes were analyzed.

The predominance of genomes originating from North America reflects the region’s leading role in clinical trials, holding 44.19% of the market in 2024 (37). More remarkable, however, is the absence of an isolation location in over a third of the data. The sharp rise in available genomes after 2006 likely reflects the introduction of the first commercially available DNA sequencing platform in 2005, which enabled large-scale sequencing efforts (38). Yet, similar to the location, the isolation date remains unknown for nearly two-fifths of the genomes. To enhance the quality and FAIR usability of sequence data, the International Sequence Database Collaboration (INSDC) introduced new standards requiring the inclusion of spatio-temporal metadata in all new ENA submissions. However, this measure has only been in place since May 2023, and although this is mandatory by now, it is still possible to report missing values (39). Same goes for the host species and the disease status (40). Public repositories are vital for advancing science by providing access to vast datasets. Nevertheless, the surge in data submissions, driven by cheaper and faster sequencing technologies, has outpaced the ability to rigorously check metadata quality (41). But without proper metadata, meaningful comparisons and global context are impossible, diminishing the value of rapidly generated sequence data. Yet, effective data sharing is essential for transparency, reproducibility, and enabling others to build on existing work. Without improved practices, the true utility of sequence data remains in question (42).

Although overall GBS colonization rates are similar worldwide, the distribution of serotypes and sequence types varies by region, host species, and clinical presentation. Two global reviews from 2023 and 2012 reported serotype III as the most prevalent, followed by serotype Ia and V (1,18). When serotype III subtypes are combined, this pattern closely mirrors the distribution identified here. Serotype III is predominantly linked to neonatal meningitis (21,43), while serotypes Ia and V are more often associated with invasive infections in non-pregnant adults (44,45). Given that over half of the genomes analyzed here originate from humans, this distribution is expected. However, it remains unclear whether the same pattern holds across non-human hosts globally. Although serotype IX was only recently identified (46), it already occurs more frequently than some other serotypes in certain regions. Increasing attention should also be given to the rising prevalence of serotype IV. Recent studies highlight this serotype as a hotspot for genetic recombination, evidenced by its occurrence across diverse genetic backgrounds (47). The growing diversity of GBS serotypes, combined with the potential for capsular switching, poses a major challenge for vaccine development, since capsular polysaccharide based vaccines may exert selective pressure that enables virulent genotypes to escape coverage. Different studies already demonstrated such switching from serotype III to IV within the hypervirulent CC17 lineage, underscoring that even highly conserved clones can alter one of their main vaccine targets (48,49). Several studies consistently report CC23, CC1 and CC19 as the most frequent lineages in global GBS collections, which indicates that these CCs represent stable and widely distributed lineages within the global GBS population (22,50,51).

Previous research found CC17 is almost exclusively linked to subtype III-2 and identified III-2/ST17 as a particularly virulent GBS clone, specifically associated with late-onset neonatal disease (50,52–55). Our investigations confirm the strong association between serotype III-2 and CC17, although no conclusions about disease status can be made due to missing metadata. Likewise, the III-1/ST19 combination is well known, and appears particularly in early-onset neonatal disease, often associated with colonizing isolates (43,52,56). The strong linkage between CC23 and serotype Ia is also well recognized, particularly in cases of invasive disease in non-pregnant adults (49,57). CC459 stood out due to its high prevalence of the macrolide resistance gene *erm(A)*, which is clinically relevant since this lineage is strongly associated with serotype IV. A 2015 study reported that ST459 strains dominate the serotype IV population responsible mainly for adult disease (58). Molecular analyses also confirm the strong link between CC1 and serotype V, with most serotype V isolates classified as ST1, while ST19 constitutes the main non-ST1 background (59,60).

Such strong serotype specificity suggests stable clonal lineages and preservation of capsule types, which can serve as useful markers for targeted surveillance. In contrast, some lineages such as CC1 and CC19 are highly heterogeneous, with CC1 showing the broadest serotype diversity. This variation has been proposed to result from an already diverse ancestral population rather than ongoing recombination (61). But most importantly, such heterogenous lineages make vaccination strategies more difficult and may harbor non-vaccine strains.

Although some studies describe serotype III as the predominant serotype in Africa, detailed data on the most common subtype remain lacking (62–64). Nevertheless, across serotype III isolates overall, III-1 and III-2 are the most frequently observed subtypes (52). Serotype III-4 and ST283 has been linked to disease in fish in several studies from Asia (13,65). Yet, more detailed metadata about the host species is needed to determine whether this represents a fish-associated lineage or possibly a case of zoonotic transmission. A study from 2021 described serotype Ib as more prevalent in Asia than in other regions of the world (66). This cannot be directly confirmed, as serotype Ib occurs at similar proportions in North America and even predominates in Australia, though the very low overall number of genomes there must be considered.

AMR is an increasingly relevant concern in GBS, impacting both treatment options and future prevention strategies. Penicillin remains the first-line treatment for colonized pregnant women, with clindamycin, erythromycin, and vancomycin recommended as alternatives in cases of penicillin allergy (67). However, resistance to second-line antibiotics like erythromycin and clindamycin has continued to rise, leading to updated treatment guidelines that now recommend cephalosporins or vancomycin as alternative therapies for adults (68). Only a small number of genomes contained genes potentially associated with beta-lactam resistance. However, since the primary mechanism for beta-lactam resistance in GBS involves amino acid substitutions in penicillin-binding proteins (PBPs) (69), this aspect was not addressed in the present analysis. Notably, MLSB resistance genes, particularly *erm(B)* and *erm(A)*, represented the second most prevalent drug class in this dataset, highlighting an important trend that should be carefully considered in future research on GBS antimicrobial resistance. Severe invasive GBS infections are often treated with a combination of penicillin and gentamicin (68). Among the genomes displaying aminoglycoside resistance, 340 specifically harbored genes conferring gentamicin resistance. In light of such emerging resistance trends, alternative antibiotic classes should be considered.

Over 80% of the analyzed genomes harbored genes for tetracycline resistance. This resistance is well known (70,71) and largely stems from the widespread use of these antibiotics since their introduction in 1948, which likely promoted the clonal expansion of resistant GBS strains. This expansion has, in turn, contributed to the emergence of lineages better adapted to human colonization and infection, most notably the dissemination of the hypervirulent ST17 clone (24,72). Tetracycline resistance genes are often located on the same mobile genetic elements as erythromycin resistance genes, especially with *erm(B)* frequently co-occurring with *tet(M)* across multiple species (71) which is also evident in this data set. While *tet(M)* is the predominant tetracycline resistance gene in GBS, mobile elements carrying other determinants like *tet(O)* or *tet(S)* have also been reported (24,73). Similarly, other erythromycin resistance determinants are frequently linked to *tet* genes and conjugative transposons capable of horizontal transfer, which can spread rapidly among GBS lineages, increasing the risk of widespread antimicrobial resistance (73). The combination of specific resistance genes should be closely monitored, as their co-occurrence may indicate the spread of multidrug-resistant lineages. The increasing spread of AMR genes in the last decades poses a serious problem and underscores the need for continuous monitoring in order to adapt prevention strategies in a targeted manner. But, the detection of AMR genes does not automatically imply phenotypic resistance. Resistance requires gene expression, and in some cases, the presence of an AMR gene may only partially reduce antibiotic susceptibility without reaching clinical resistance thresholds. Conversely, isolates may gain or lose resistance through other mutations. Therefore, genotypic results should be interpreted with caution and ideally supported by phenotypic testing. Moreover, standardized and comprehensive metadata, such as isolation source and disease type, are essential to enable accurate interpretation of resistance trends and to facilitate global comparisons.

Distinct clonal complex/serotype combinations are associated with characteristic gene repertoires and span a continuum from flexibility to niche-specific specialization. These differences likely contribute to lineage-specific invasive potential across host groups, exemplified by the hypervirulence of CC17/III-2 in neonates and the comparatively low neonatal invasiveness of CC23/Ia and CC459/IV and the tendency to cause disease in adults. All identified gene patterns point to distinct evolutionary and ecological strategies among the groups. The enrichment of mobile genetic elements and phage-associated genes suggests an enhanced genomic plasticity, likely reflecting frequent exposure to horizontal gene transfer. By contrast, II/22 shows a focused enrichment of siderophore and metal-uptake genes, indicating specialization for iron-limited host niches, while III-2/17 accumulates virulence and adhesion genes, especially the accessory SecA2/SecY2 system, pointing to enhanced host interaction and invasiveness. Beyond the canonical secretion (Sec) pathway, GBS carries an accessory SecA2/SecY2 secretion system. This alternative locus is apparently restricted to the hypervirulent ST17 lineage that encodes the related Srr2 adhesin. Srr2 is specific for ST17 strains and mediates high-affinity binding to fibrinogen and plasminogen, thereby enhancing adhesion, resistance to phagocytic killing and persistence in a murine meningitis model (74). The presence of these genes in group III-2/17 supports the reliability of our analysis. Together, these results indicate that even a small set of accessory genes may strongly influence the adaptation of GBS isolates.

## Conclusion

The aim of this study was to assess the overall state of research on GBS. However, due to substantial gaps in the available metadata, a comprehensive evaluation is not possible. For instance, data from camelids seems entirely missing, and for human-derived isolates it is mostly unclear whether they originate from adult or neonatal cases. Large-scale comparative genomics, especially studies dissecting genetic variation and adaptive genome dynamics, provides key insights into multi-host pathogens like GBS. But the continuous generation of sequencing data without adequately structured and curated metadata limits their utility. The lack of standardized and comprehensive metadata represents a major obstacle in modern genomics and personalized medicine, restricting both biological insight and meaningful medical application. Safeguarding the integrity of data and metadata is not optional, it is essential, and compromising on data quality ultimately compromises science itself.

## Notes

### Competing Interest Statement

The authors have declared no competing interest.

