## Supplementary Figures 1+2 for "Towards a holistic epidemiology of *Streptococcus agalactiae* using the BakRep repository"

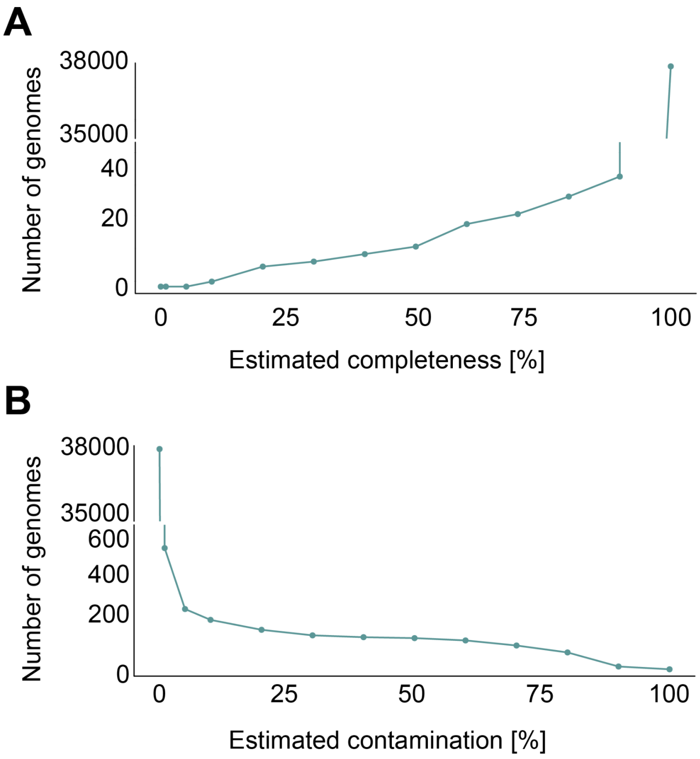


**Supplemental Figure 1:** Distribution of completeness **(A)** and contamination **(B)** level of all GBS genomes contained in BakRep v2.

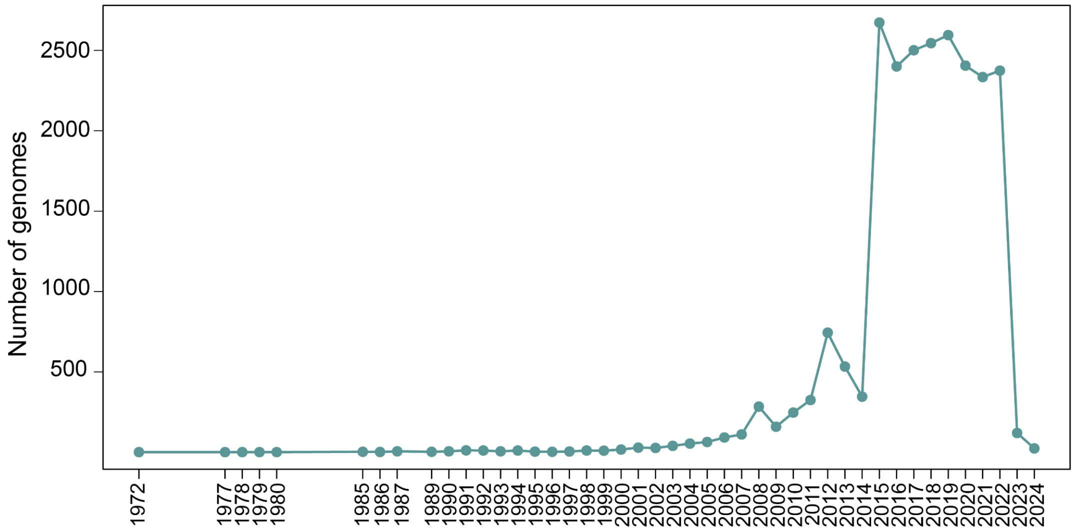


**Supplemental Figure 2:** The yearly rate of GBS genomes contained in BakRep v2. The linear chart shows the number of genomes per year from 1972 to 2024.
